# Individual variation in meiotic crossover positioning, rate and interference are associated with distinct genetic processes in domestic pigs

**DOI:** 10.1101/2024.06.20.599484

**Authors:** Cathrine Brekke, Arne B. Gjuvsland, Peer Berg, Susan E. Johnston

**Author notes:** Department of Animal and Aquacultural Sciences, Norwegian University of Life Sciences, Oluf Thesens vei 6, 1433 Ås, Norway (Cathrine Brekke). Corresponding author: Cathrine Brekke.

## Abstract

Meiotic crossovers are essential for proper chromosome segregation, and provide an important mechanism for adaptation through linking beneficial alleles and purging deleterious mutations. However, crossovers can also break apart beneficial alleles and are themselves a source of new mutations within the genome. The rate and distribution of crossovers shows huge variation both within and between chromosomes, individuals and species, yet the molecular and evolutionary causes and consequences of this variation remain poorly understood. A key step in understanding this variation is to understand the genetic architecture of how many crossovers occur, where they occur, and how they interfere, as this allows us to identify the degree to which these factors are governed by common or distinct genetic processes. Here, we investigate individual variation in crossover count, crossover interference (ν), and crossover positioning measured as both intra-chromosomal allelic shuffling and distance to telomere (Mb), in a large genotyped breeding population of domestic pigs. Using measures from 82,474 gametes from 4,704 mothers and 271 fathers, we show that crossover traits are heritable within each sex (h^2^ = 0.03 - 0.11), with the exception of male crossover interference. Crossover count and interference have a strongly shared genetic architecture in females, mostly driven by variants at *RNF212*. Female crossover positioning is mediated by variants at *MEI4*, *PRDM9*, and *SYCP2*. We also identify tentative associations at genomic regions corresponding to *CTCF* and *REC114/REC8/CCNB1IP1* (crossover count), and *ZCWPW1* and *ZCWPW2* (crossover positioning). Our results show that crossover count and crossover positioning in female pigs have the capacity to evolve somewhat independently in our dataset.

## Introduction

Meiotic recombination through chromosomal crossovers is a fundamental feature of sexually-reproducing species. In many species, it is required for the proper segregation of chromosomes into eggs and sperm, with most species having a minimum requirement of one crossover per chromosome pair [1]. It also facilitates adaptation by bringing together beneficial alleles at linked loci [2–4], and providing a mechanism to purge deleterious alleles from genomes [5,6]. However, the crossover process also has risks, as it increases the rate of mutations at double strand break (DSB) repair sites [7,8] and can uncouple beneficial alleles at linked loci [9]. The genetic mechanisms of recombination and crossing-over are highly conserved across eukaryotes, yet there is a high diversity in their sequence and function [10,11]. Furthermore, the rate and distribution of crossovers show a huge diversity both within and between chromosomes, individuals, sexes, populations and species [12,13]. A key step in understanding this diversity is to determine its genetic architecture at the individual level (i.e., the underlying gene variants and their relative effects on crossover processes), as this can help identify the molecular mechanisms underpinning variation, its evolutionary capacity and constraints, and potential effects on downstream evolutionary processes [12,14,15].

To gain a full picture of the causes and consequences of crossover variation, there are several non-independent processes that must be considered. First, we must consider ***how many crossovers occur***, as fewer crossovers may increase the risk of non-disjunction, whereas too many may lead to genome instability and increased rates of deleterious mutation [1,16,17]. The genetic basis of crossover count has been investigated in several mammal species, including humans, cattle, pigs, sheep, and deer [18–26]. A number of large effect loci have been identified, including *RNF212*, *RNF212B*, *MEI1*, *MSH4*, *PRDM9*, and *REC8*, among others, with their functions associated with meiotic processes such as crossover designation and DSB initiation and repair [10]. In particular, the locus *RNF212* and/or its paralogue *RNF212B* are consistently associated with crossover count variation in almost all of these studies, and likely exhibits a dosage dependent effect on crossover number [27].

Second, we must consider ***where crossovers occur***. The positioning of crossovers can influence the proportion of alleles on a chromosome that will be coupled or uncoupled (or “shuffled”). For example, a crossover situated at the end of a chromosome pair will shuffle a relatively small proportion of alleles as most linked variants will remain intact, whereas a crossover situated in the centre of a chromosome pair will shuffle a relatively high proportion of alleles, as all loci on one side of the crossover are uncoupled with those on the other side [28]. This distinction is likely to have evolutionary consequences in terms of the rate of generation of novel linked allelic variation within populations [28]. Patterns of crossover positioning are likely to be affected by the physical structure of meiotic chromosomes. During meiosis, the DNA is structured into chromatin loops tethered along an axis structure, and held in close proximity by a protein structure called the synaptonemal complex (SC) [30], and shorter axes/SCs and longer DNA loops are associated with reduced crossover rates [13,31–33]. Similarly, crossovers can vary relative to local chromatin accessibility, gene promotor regions, methylation patterns, structural variants, and proximity to centromeres and telomeres [34,35]. Less is known about individual variation in crossover positioning in mammals, although heritable variation has been identified in house sparrows and Atlantic salmon [36–38] and changes in crossover landscapes have been observed under domestication in tomatoes, rye and barley [33,39,40]. Fine-scale crossover landscapes in mammals are also often mediated by the rapidly evolving locus *PRDM9*, which binds to particular allele-specific DNA sequence motifs and promotes DSB formation in recombination “hotspots” of around 1-10kb in width [41–43].

Third, we must consider ***how crossovers interfere*** with one other. The distribution of crossovers along a chromosome is affected by the phenomenon of “crossover interference”, where a crossover forming in one position will reduce the probability that more crossovers will form nearby [44–47]. Several mechanisms have been proposed to explain how and why crossover interference is pervasive across eukaryotes, relating to mechanical stress on the chromosome, telomere initiation of crossovers, and aggregation of pro-crossover proteins into coarse clusters (“coarsening model”), among others [30,42]. Crossover interference may also be affected by chromosome length and structure, where shorter chromosomes, axes/SCs and larger DNA loops may increase the effects of interference at the base-pair scale [48]. Heritable variation in individual crossover interference has been identified in cattle, with some variance explained by the locus *NEK9* [48]. However, such studies remain rare, as analyses rely on information from two or more crossovers on the same chromosome which are rare at the genome-wide scale, meaning that large sample sizes per individual are required to measure this metric with accuracy [46]. In *Arabidopsis thaliana*, the locus *Hei10* (in the same E3 ubiquitin ligase family and with a similar function to *RNF212*) has been associated with crossover interference through dosage along the chromosome, corresponding to the coarsening model described above [49,50].

All three of these processes interact. Crossover interference will affect how many crossovers can be placed on the chromosome; the number of crossovers will also affect levels of allelic shuffling; and crossover positioning will determine if there is enough distance for other crossovers to overcome interference. Yet, it remains unclear the degree to which these three phenomena are phenotypically and genetically correlated, and in turn, if these processes have the capacity to evolve independently. Understanding these distinctions may have importance in separating the mechanistic and evolutionary drivers of recombination e.g. through ensuring fertility through obligate crossing over, or through novel haplotype generation/haplotype conservation via crossover positioning. We must also consider that crossover processes are occurring within two distinct gametic environments: oogenesis and spermatogenesis. Differences in crossover rates and distribution between the sexes is a near universal phenomenon in eukaryotes (known as “heterochiasmy”, or “achiasmy” when recombination is absent in one sex; [18,24,34,35,51,52]), and there is increasing evidence in vertebrates that the genetic architecture of crossover processes is also different between the sexes [18,24,34,35,51,52]. Therefore, determining phenotypic and genetic variation and correlations in these processes, in both males and females, will allow us to have a better understanding of the molecular mechanisms, evolutionary potential, and evolutionary constraints operating on these processes [42].

Domestic pigs represent an excellent system to explore variation in crossover processes, as large litters allow for the quantification of crossover counts, crossover positioning, and crossover interference in both males and females. Previous studies have shown that crossover count in pigs is heritable, with a polygenic architecture in males and an oligogenic architecture in females associated with variants at *RNF212*, *SYCP2*, *PRDM9*, *MEI1* and *MSH4* [53]. However, the genetic basis of crossover positioning and crossover interference, and their relationship with crossover count, remains unknown. In this study, we integrate SNP data with a large breeding pedigree of Large White pigs to quantify crossover positions in 82,474 gametes transmitted from 4,704 mothers and 271 fathers. We quantify crossover count, crossover positioning, and crossover interference, and determine the heritability, genetic correlations, and genetic architecture of these traits within each sex. We show that crossover count and interference show a shared genetic architecture in females, mostly driven by variants at *RNF212*, whereas crossover positioning is mainly mediated by variants at *MEI4*, *PRDM9*, and *SYCP2*.

## Results

### Crossover phenotype dataset

We used data from a breeding pedigree of Large White pigs genotyped at 50,705 SNP markers to characterise autosomal crossover positions in gametes transmitted from focal individuals to their offspring. SNP positions are known relative to the Sscrofa11.1 reference genome assembly. Our dataset comprised of 41,237 gametes transmitted from 4,704 unique females, and 41,237 gametes transmitted from 271 unique males. We used the crossover positions to estimate the four crossover phenotypes across all autosomes per gamete:

- ***Crossover Count***: the total number of crossovers within a gamete.
- ***Intra-Chromosomal Allelic Shuffling (Crossover positioning***, 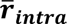): the probability that two randomly chosen loci on the same chromosome are uncoupled within a gamete. This metric quantifies *crossover positioning* i.e. higher values are indicative of chromosomes having crossovers closer to the centre, whereas lower values are indicative of crossovers occurring towards chromosome ends [28].
- ***Distance to Telomere (Mb)***: the mean distance between the telomere and the closest crossover on the same chromosome arm within a gamete.
- ***Crossover Interference (*ν)**: the strength of crossover interference in all gametes transmitted from each individual, as defined by the shape parameter ν. This was fit with a Houseworth-Stahl interference escape model, which models both interfering (Class I) and non-interfering (Class II) crossovers [26,40,54].

All phenotypes showed differences in their mean and/or distribution between females and males, with the exception of intra-chromosomal shuffling (Table 1, Figure 1). Crossover counts were higher in females, but the distance to telomere and strength of crossover interference were higher in males (Table 1, Figure 1). Crossover count and intra-chromosomal shuffling were positively correlated (Pearson’s r = 0.581, P < 0.001), as were crossover count and the distance to telomere (r = 0.130, P < 0.001), and intra-chromosomal shuffling with distance to telomere (r = 0.463, P < 0.001). Crossover interference was weakly negatively correlated with all other crossover phenotypes in females (r = -0.155 to -0.071; P < 0.001) and was not correlated with other phenotypes in males (P > 0.05, Figure 1).

**Figure 1.**
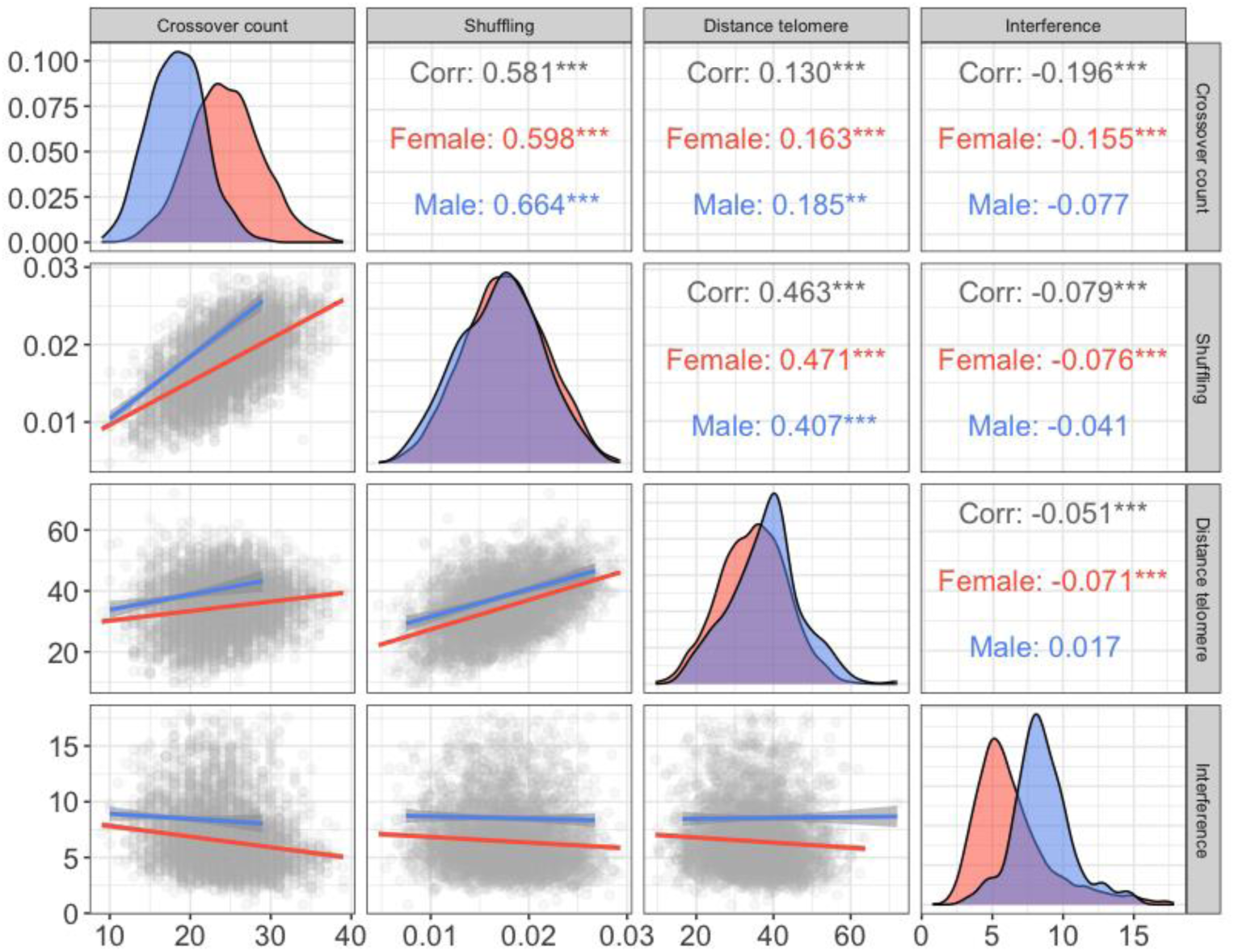
Sex-specific distributions of crossover phenotypes (diagonal panels) and their sex-averaged and sex-specific linear regressions (lower triangle) and Pearsons correlations (upper triangle). Male and female phenotypic correlations and distributions are indicated in blue and red, respectively. *** indicates a correlation significance of P < 0.001. The shaded area around the regression lines indicates the 95% confidence interval.

**Table 1.** Sample sizes, means and estimated variance components for all crossover phenotypes for males (M) and females (F). N_FIDs_ is the number of unique focal individuals, N_obs_ is the total number of gametes (i.e. measures), V_P_ is the phenotypic variance, h^2^ is the narrow-sense heritability, pe^2^ is the proportion of permanent environmental variance, r_A_ is the genetic correlations between males and females. Values in parenthesis are the standard errors.

| Crossover Phenotype | Sex | $N_{\text{FIDs}}$ | $N_{\text{obs}}$ | Mean | $V_p$ | $h^2$ | $pe^2$ | Cross-sex $r_A$ |
| --- | --- | --- | --- | --- | --- | --- | --- | --- |
| Crossover Count | M | 271 | 41,237 | 18.20<br>(3.37) | 11.43<br>(0.11) | 0.06<br>(0.01) | $0.43e^{-10}$<br>($0.42e^{-12}$) | 0.48<br>(0.10) |
| | F | 4,704 | 41,237 | 24.37<br>(4.58) | 21.04<br>(0.17) | 0.11<br>(0.01) | $0.11e^{-3}$<br>(0.02) | |
| Intra-Chr Allelic Shuffling ($\bar{r}_{\text{intra}}$) | M | 271 | 41,237 | 0.0172<br>(0.004) | $0.16e^{-4}$<br>($0.14e^{-6}$) | 0.04<br>(0.01) | $0.40e^{-2}$<br>(0.02) | 0.26<br>(0.14) |
| | F | 4,704 | 41,237 | 0.0175<br>(0.004) | $0.18e^{-4}$<br>($0.14e^{-6}$) | 0.09<br>(0.01) | 0.02<br>(0.02) | |
| Distance to Telomere (Mb) | M | 271 | 41,237 | 37.25<br>(9.48) | 90.19<br>(0.75) | 0.03<br>(0.01) | 0.01<br>(0.01) | 0.18<br>(0.14) |
|  | F | 4,704 | 41,237 | 34.80<br>(8.65) | 74.90<br>(0.57) | 0.07<br>(0.01) | 0.01<br>(0.01) |  |
| Crossover Interference (v) | M | 271 | 271 | 8.54<br>(2.07) | 12.76<br>(1.10) | 0.00<br>(0.00) | * | ** |
|  | F | 4,704 | 4,704 | 6.45<br>(2.77) | 24.24<br>(0.70) | 0.08<br>(0.02) | * |  |

\*this was not estimated as there is only one measure of crossover interference per individual.
\*\*model did not converge.

### Heritabilities and Genome-wide Association Studies (GWAS)

All crossover phenotypes were heritable within each sex (h^2^ = 0.03 - 0.11), with the exception of male crossover interference, which was not significantly heritable (Table 1). Females consistently had higher heritabilities than males, and had higher phenotypic variance for all traits, except the distance to telomere which had higher phenotypic variance in males (Table 1). The cross-sex genetic correlations were positive for each crossover phenotype, but were low to moderate in magnitude, ranging from 0.18 for distance to telomere to 0.48 for crossover count (Table 1). Crossover count, intra-chromosomal allelic shuffling and distance to telomere had significant positive additive genetic correlations within each sex (r_A_ = 0.36 - 0.86), whereas crossover interference in females had negative additive genetic correlations with all other crossover phenotypes (r_A_ = -0.32 to -0.97; Table 2). We were unable to estimate additive genetic correlations between male crossover interference and other crossover traits, as models failed to converge given that this trait was not heritable.

**Table 2.** Additive genetic correlations of crossover phenotypes within each sex. Female correlations are shown in the upper triangle and male correlations are shown in the lower triangle. Values in parentheses are the standard errors.

| Crossover Trait | Crossover count | Intra-Chr. Allelic Shuffling | Distance to telomere | Crossover Interference |
| --- | --- | --- | --- | --- |
| Crossover count | - | 0.70 (0.03) | 0.37 (0.05) | -0.97 (0.05) |
| Intra-Chr. Allelic Shuffling | 0.64 (0.05) | - | 0.86 (0.02) | -0.59 (0.11) |
| Distance to telomere | 0.36 (0.08) | 0.79 (0.06) | - | -0.32 (0.13) |
| Crossover Interference | * | * | * | - |

\*model did not converge.

Genome-wide association studies were carried out using a larger imputed dataset of 524,587 SNPs, and were run within each sex separately. All crossover phenotypes had significant associations in the GWAS studies in females, whereas there were no significant associations in males (Figure 2, Table 3). For all significant associations, we identified candidate genes in the proximity of the most highly associated SNPs based on gene ontology (GO) terms associated with meiotic processes. All significant SNP associations are provided in Table S1. Individual plots of significant regions are provided in Figure S1. All candidate gene positions are provided in Table S2, with associated GO terms provided in Table S3. We describe results for each phenotype in detail:

**Figure 2.**
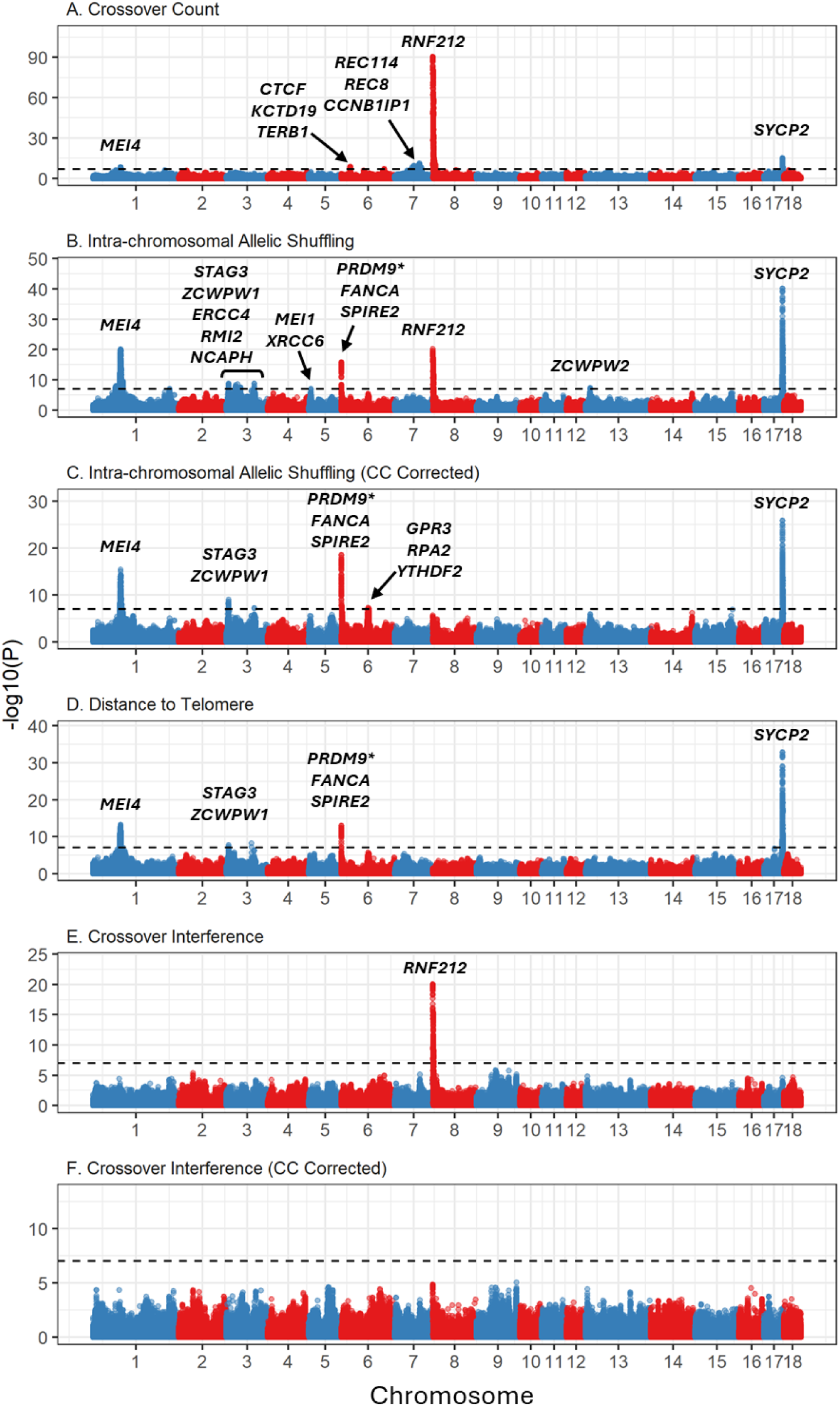
Genome-wide association plots for 6 crossover traits (Panels A-F) in females. Associations are displayed at 554,768 SNPs. Sample sizes are provided in Table 1. CC Corrected indicates that crossover count was fitted as a fixed covariate in the model. Gene names above significant peaks indicate direct candidate loci. The dashed line indicates the genome-wide significance threshold at α= 0.05. Association statistics have been corrected with the genomic control parameter λ. Information on significant loci and candidate genes are provided in Table 2 and Tables S1-3. Plots of significant regions are provided in Figure S1. \**PRMD9* is annotated as *PRDM7*; see main text for discussion.

**Table 3.**
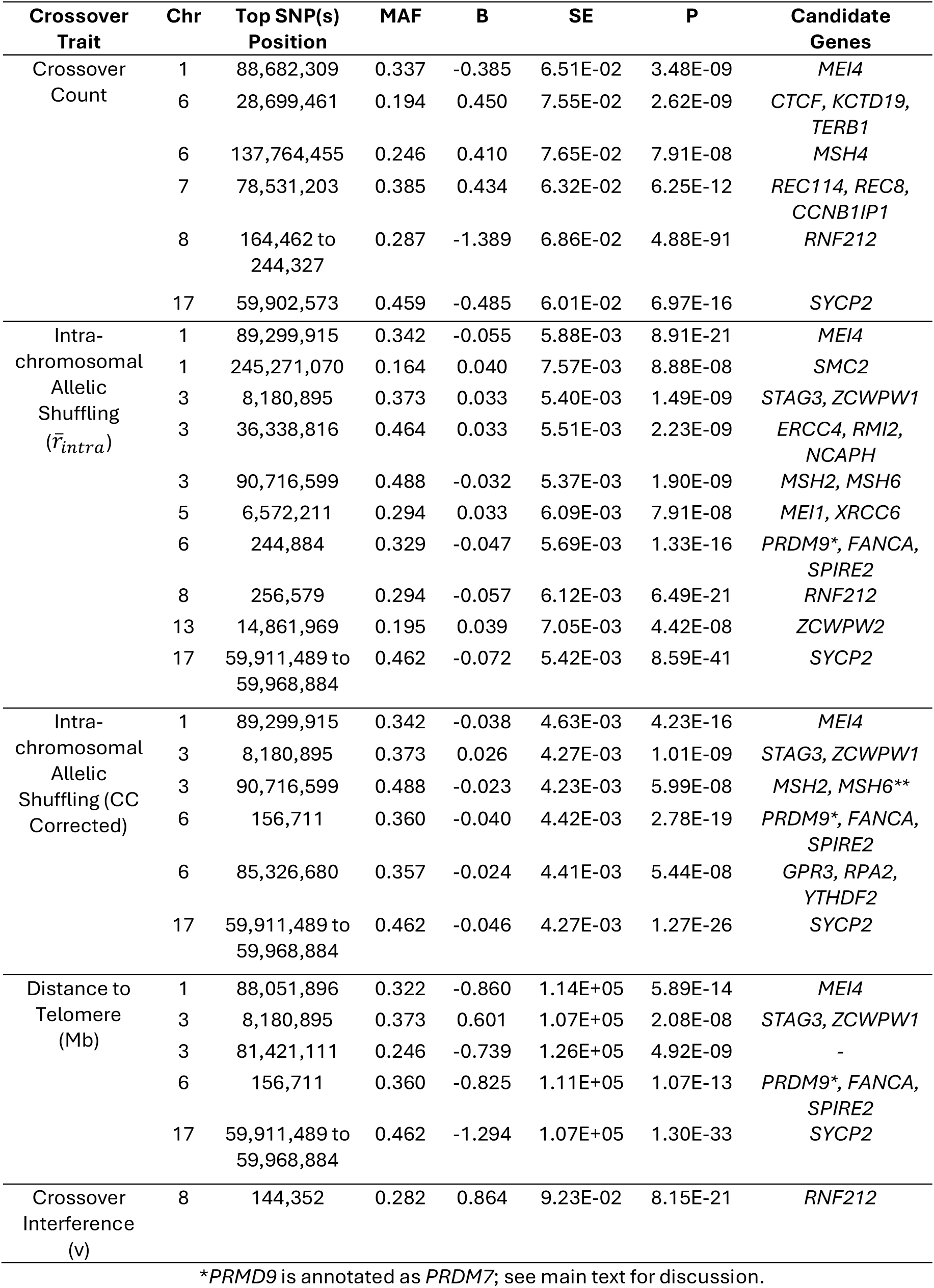
Genomic regions associated with crossover phenotypes in females. This table includes information for the most highly associated SNP loci; full results are provided in Table S1. MAF is the minor allele frequency. B is the effect size of the minor allele with standard error SE. P is the association significance after correction for genomic control. Candidate genes are those directly associated with meiotic processes. Full gene names, positions, and associated Gene ontology terms are provided in Tables S2 and S3.

#### Crossover Count

This trait has been investigated previously in the same dataset [42], but was reanalysed here with a leaving-one-chromosome-out GWAS model, which identified new associations. Crossover count was 1.34 times higher in females than in males and was heritable in both sexes (h^2^ = 0.06 and 0.11 in males and females, respectively), with a moderate cross-sex genetic correlation (r_A_ = 0.48; Table 1). A GWAS identified six regions on five chromosomes that were significantly associated with female crossover count, corresponding to ten candidate genes. The strongest association was observed at *RNF212* on chromosome 8 (Figure 2A, Figure S2, Table 3). Significant associations were also identified at two regions in common with intra-chromosomal allelic shuffling and distance to telomere, and corresponded to *MEI4* and *SYCP2* on chromosomes 1 and 17, respectively. There were three regions uniquely associated with crossover count, including a region on chromosome 6 corresponding to the candidate genes *CTCF*, *KCTD19* and *TERB1*, and a region on chromosome 7 corresponding to the candidate genes *REC114*, *REC8* and *CCNB1IP1*.

#### Intra-chromosomal Allelic Shuffling (Crossover positioning)

This trait was heritable in both males and females (h^2^ = 0.04 and 0.09, respectively), but the means were not significantly different between the sexes (Table 1). Despite this, the cross-sex additive genetic correlation was relatively low (r_A_ = 0.26, Table 1), indicating the genetic architecture of this trait is largely independent between the sexes. Within each sex, this trait was strongly correlated with crossover count (0.64 and 0.7 in males and females, respectively) and distance to telomere (0.79 and 0.86 in males and females, respectively; Table 2). GWAS identified significant associations in ten regions on seven chromosomes, corresponding to 18 potential candidate genes, including *RNF212* (Figure 2B, Table 3). As intra-chromosomal allelic shuffling is correlated with crossover count, we ran an additional GWAS including crossover count fit as a fixed covariate in the model (Figure 2C, Table 3). After this correction, only six regions on 4 chromosomes remained significant, and the association with *RNF212* was no longer significant. The strongest association was observed at *SYCP2* on chromosome 17, with additional strong associations shown at *MEI4* on chromosome 1, and *PRDM9*, *FANCA* and *SPIRE2* on chromosome 6. Note that *PRDM9* on the pig genome is annotated as *PRDM7*; however, we hereafter refer to this locus as *PRDM9*, based on identification and discussion in previous publications, and high conservation of sequence and function between *PRDM7* and *PRDM9* [18,21,22,25,26,51].

#### Distance to Telomere

This trait was heritable in both males and females (h^2^ = 0.03 and 0.07, respectively; Table 1). Males tended to have crossovers further from telomeres than females, although this difference was not significant; nevertheless, the cross-sex additive genetic correlation was relatively low (r_A_ = 0.18, Table 1, Figure 1). This trait was highly correlated with intra-chromosomal shuffling in each sex (r_A_ = 0.79 and 0.86 in males and females, respectively), but less so with crossover count (r_A_ = 0.36 and 0.37 in males and females, respectively; Table 2). GWAS identified significant associations in five regions on 4 chromosomes (Figure 2D, Table 3). Similarly to intra-chromosomal allelic shuffling, the strongest association was observed at *SYCP2* on chromosome 17, with additional strong associations shown at *MEI4* on chromosome 1, and *PRDM9*, *FANCA* and *SPIRE2* on chromosome 6.

#### Crossover interference

Both males and females showed positive crossover interference, i.e., crossovers are more distantly spaced than expected by chance; interference was stronger in males than in females (Table 1). The level of interference was similar to other mammal systems, such as humans, cattle, sheep, and mouse, which range from ν = 6.7 in female cattle to ν = 11.65 in male mouse [18,21,22,25,26,51]. A small proportion of crossovers were designated as non-interfering (i.e., Class II crossovers) although this was not significantly different from zero in either sex (4.9% and 4.3% of crossovers in males and females respectively, s.e. = 6%). Crossover interference was only heritable in females (h^2^ = 0.08, Table 1) and showed a very strong negative additive genetic correlation with crossover count (r_A_ = -0.97, Table 2). It should be noted that non-significant results in males may be due to the relatively small sample size for this metric (N = 271 in males vs 4,704 in females). Only one region was significantly associated with female crossover interference, corresponding to *RNF212* on chromosome 8 (Table 2, Figure 2E). However, this association was no longer significant when fitting crossover count as a fixed covariate in the model (Figure 2F), indicating that *RNF212* has a strong effect on both crossover count and crossover interference in females of this population.

## Discussion

This study has shown that individual variation in crossover count, positioning, and interference is heritable and differs between the sexes in a large breeding population of domestic pigs. Heritabilities were generally low in both sexes (h^2^ < 0.1), but large numbers of meioses sampled from thousands of individuals allowed us to identify 14 distinct genomic regions associated with crossover trait variation in females. Crossover rate and interference were strongly negatively correlated and associated with a large effect locus corresponding to *RNF212*. Crossover positioning was partly correlated with crossover rate, and was associated with large effect loci corresponding to *MEI4*, *PRDM9* and *SYCP2*. Crossover traits in males had lower heritabilities and no genomic regions with significantly large effects on the traits; while this may be reflective of polygenic architectures, it may also be that the small number of unique males in this data set (271 males compared to 4,704 females) led to reduced power to identify moderate to large effect loci. Here, we discuss in more detail the role and function of candidate loci, variation in crossover interference, why genetic variation is maintained and potential breeding outcomes, and future directions for research.

### *RNF212* as a locus for crossover count and interference

Crossover count and interference showed strong associations with the locus *RNF212*. This locus and/or its paralogue *RNF212B* are consistently associated with crossover count in nearly every study in mammals published to date [45,46,55]. Functional studies in e.g. mice have shown that RNF212 proteins localises to recombination sites early in the crossover designation process, and is a dosage sensitive regulator of crossover formation [45,46]. The genetic correlation between crossover count and interference was strongly negative, and correcting interference for crossover count removed all association with the *RNF212* loci. Therefore, another plausible mechanism is that lower crossover counts are mediated by higher crossover interference. This is suggestive of the coarsening model of crossover interference observed in *Arabidopsis thaliana* mediated by Hei10, a protein in the same family of E4 ligases which exhibit a conserved behaviour during meiosis [56–59]. In *A. thaliana*, a number of small Hei10 foci form on chromosomes during synapsis, which in turn aggregate into fewer, larger, distinct foci along the chromosome (i.e. they undergo a coarsening process), which correspond to final crossover sites [60]. Whilst we cannot directly infer the mechanism by which *RNF212* affects these processes in the pigs, our findings highlight a likely inter-dependence of crossover count and the strength of crossover interference in this population.

### Additional candidate loci associated with crossover count

Two additional genomic regions were associated with crossover count. One contained the locus *CTCF*, a DNA binding protein that is associated with the anchoring of chromatin loops and the boundaries of topologically associated domains [56–59]. There is evidence that CTCF contributes to the DNA loop and axis organisation of chromosomes during meiosis, and may interact with PRDM9 binding sites [61,62]. Another region contained the meiotic recombination proteins *REC8*, *CCNB1IP1* and *REC114*; this region has been association with crossover count in humans, deer and cattle [63]. *CCNB1IP1* is the orthologue of Hei10, and all three loci are required for the crossover process [64,65], and there is evidence that *REC8* can be enriched at CTCF sites in mouse spermatocytes [66].

### *SYCP2*, *MEI4* and *PRDM9* as loci for broad-scale crossover positioning

This study showed a strong association of the loci *MEI4* and *SYCP2* on crossover positioning, through intra-chromosomal shuffling and distance to the telomere. SYCP2 is an important component of the synaptonemal complex that is required for synapsis and recombination during meiosis and functional studies in zebrafish has shown that SYCP2 is associated with synaptonemal complex assembly initiation close to telomeres [66,67]. In mice, association of MEI4 to the chromosome axes is a requirement for meiotic DSB formation, and can be a limiting factor in this regard [68], and there is evidence that MEI4 is directed to DSB sites by the PRDM9 protein [69,70]. MEI4 also forms a complex with REC114 (see above), which is essential for meiotic DSB formation [39]. *SYCP2* and *MEI4* were also shown to have a weaker but significant effect on crossover count, indicating that crossover positioning may also influence the number of crossovers that can occur. The association between *PRDM9* and broad-scale positioning is less clear, as this locus is generally considered as a mediator of the positioning of fine-scale recombination hotspots (i.e. in regions of 1-2kb) [71]. Different alleles of the *PRDM9* zinc-finger array bind to different sequence motifs throughout the genome; therefore, this locus could feasibly affect broad-scale positioning if there is a difference in motif abundances associated with broad-scale features of the genome [72]. In addition, the interaction of this locus with MEI4 may lead to broad-scale effects on top of its role in fine-scale variation. Interestingly, crossover positioning also showed weak but significant effects with the loci *ZCWPW1* and *ZCWPW2*. *ZCWPW1* has co-evolved with *PRDM9*, and ZCWPW1 is recruited to recombination hotspots by PRDM9, and is required for DSB repair during meiosis [73]. In addition, in species where *PRDM9* has been lost, *ZCWPW1* and *ZCWPW2* are also more likely to be lost, suggesting a key role of both loci in the recombination process [48,74–76]. The region containing *ZCWPW1* also contained *STAG3*, a locus which is associated with all meiotic cohesion complexes and is required for normal axial localisation, DNA repair, and crossing over [53].

### Variation and sex differences in crossover interference

Our dataset had large numbers of offspring for both male and female individuals, which allowed us to conduct an analysis of crossover interference within both sexes. We found that the strength of crossover interference was higher in male pigs, which also had the lower crossover rate; this pattern has also been found in humans, cattle, dogs and mice [77]. Crossover interference was modelled using a Houseworth-Stahl model of interference [78,79], which allows both interfering and non-interfering crossovers to occur. Our results show that the number of crossovers escaping interference was not significantly different from zero. However, our crossover dataset may have reduced power to detect non-interfering crossovers, as (a) the median marker density is around 1 SNP per 31.8kb, and (b) the software used to estimated crossover positions (Lep-MAP3) uses a likelihood-based model to call crossovers based on phase changes, and may not call very short crossovers with marginal changes in map likelihoods [30,80]. Our analysis used sex specific genetic map distances rather than base pair distances to calculate crossover interference parameters. This is because it is well established that crossover interference operates on the length of the chromosome axis, i.e. physical length of the chromosome during pachytene [81,82], and the synaptonemal complex length is different in male and female meiosis and strongly correlates with genetic length in cM in both sexes [83].

### Why is genetic variation for crossover processes maintained in pigs?

Our findings add to a growing body of evidence of genetic variation underpinning crossover traits in domesticated mammal species [20,21,23,24,26,51]. Some models predict that strong selection (particularly in smaller breeding populations) should favour alleles that increase rates of recombination [81], which should in turn reduce genetic variation in recombination rates. The heritability of the crossover traits measured here are low, but there is still the evolutionary potential to modify crossover traits to reach an adaptive optimum. In addition, there is the capacity for independent evolution of crossover positioning and crossover rate, whereas interference and crossover rate are strongly genetically coupled in this population. Nevertheless, selection on crossover traits for their advantages related to breeding outcomes (i.e. through faster responses to selection) is likely to be weak. In a species such as pigs, where there are a large number of chromosomes (2N = 36), intra-chromosomal allelic shuffling contributes to a negligible amount of shuffling at the genome-wide level when compared to the independent assortment of chromosomes [28]. In addition, selection on both crossover count and crossover distribution to their maximum heritable capacity will only result in small breeding gains, as shown on simulated quantitative traits [82,83]. It is likely that the presence of crossovers provides enough of a mechanism to purge deleterious mutations and to overcome Hill-Robertson interference in the face of genetic drift [81,84].

### Conclusions

Our findings show that crossover rate, crossover positioning, and crossover interference can be driven by distinct genetic processes in domestic pigs, with some degree of shared genetic architecture. Future progress will rely on understanding the molecular mechanisms of this variation in more detail, particularly in terms of wether individual differences in chromosome architectures are the driver or consequence of genetic differences at the loci identified above. In addition, it remains to be known whether individual phenotypic and genetic variation in crossover traits are associated with individual fertility e.g. through proper segregation of chromosomes and maintenance of genomic integrity through reduced mutation rates associated with lower levels of DSB formation and repair.

## Methods

### Study population and genotype dataset

This study used genomic and pedigree data in a Large White breeding population of 95,613 individuals provided by Topigs Norsvin. All study individuals were genotyped on one of two SNP arrays: Illumina GeneSeek Custom 80K SNP chip, and Illumina GeneSeek Custom 50K SNP, with 50,705 SNPs in common between the two arrays (hereafter referred to as the 50K dataset). Quality control removed SNPs with a minor allele frequency < 0.01, genotype call rate < 0.95, and strong deviation from Hardy Weinberg equilibrium (χ_1_^2^ > 600). The physical positions of the markers were determined relative to the Sscrofa11.1 reference genome assembly. Key individuals in the pedigree were also genotyped on an Axiom porcine 660K Affymetrix array, and genotypes of all focal individuals in this study were imputed to 660K SNPs using the Fimpute genotype imputation software V2.2 [85]. Quality control removed imputed SNPs with a minor allele frequency of < 0.01, with a final dataset of 524,587 SNPs. Note that the imputed dataset was only used in the genome wide association analysis and not to infer crossovers.

### Linkage mapping and estimation of meiotic crossover positions

Individual crossover positions were estimated using the same approach as [53] using the 50K dataset. Briefly, the pedigree was ordered into three generation full-sib families, consisting of two focal individuals (female and male), their parents, and their offspring. This allowed phasing of the 50K marker dataset in focal IDs and their offspring, and consequently the identification of crossovers positions in gametes transmitted from the focal ID to their offspring. Characteristics of the crossover positions are then assigned as a phenotype of the focal ID, as this is where the meiosis took place. The software Lep-MAP3 [42] was used to construct sex-specific linkage maps at the population level, and to estimate autosomal crossover positions within individual phased gametes. Marker orders were assumed to be the same as Sscrofa11.1, and linkage maps were constructed in centiMorgans (cM) using the Morgan mapping function. Sample sizes are provided in Table 1.

### Estimation of crossover phenotypes

Four crossover phenotypes were estimated for each gamete for each focal ID (unless otherwise stated) using information on the meiotic crossover positions on autosomes. Each phenotype was determined in autosomes only to allow direct comparisons of rates between females and males. These were as follows:

#### Crossover Count

This measure is the total crossover count per gamete across all autosomes. Analysis of crossover count has previously been published for the same dataset in this breed [30,85].

#### Intra-Chromosomal Allelic Shuffling (Crossover positioning)

Intra-chromosomal shuffling, 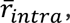 was calculated per gamete per focal ID as the probability that a pair of loci on the same chromosome are uncoupled due to a crossover event, using the following equation adapted from [28]:

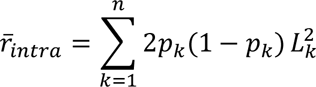

where *k* is the autosome number 1–18, n is the number of autosomes, *p* is the proportion of paternally inherited alleles, *1-p* is the proportion of maternally inherited alleles, and *L* is the length of the chromosome as a fraction of the total length of the genome.

#### Distance from telomere

To obtain individual measures of crossover distance from telomere, we measured the physical distance between a crossover event and the closest telomeric end of the chromosome in Megabase pairs (Mb). On metacentric chromosomes, the distance from telomere for a crossover on the *p* arm was the position of the crossover in Mb, and distance from telomere for a crossover on the *q* arm was the total length of the chromosome minus the position of the crossover. If a chromosome arm had more than one crossover, only the crossover closest to the telomere was counted, i.e. a maximum 2 for metacentric chromosomes (1-12) and 1 for acrocentric chromosomes (13-18). For each gamete per focal ID, we took the mean distance from a crossover to the telomere across chromosomes. Chromosomes with zero crossovers were not counted. Centromere positions were obtained from [86].

#### Crossover interference

Individual strength of crossover interference (ν) was estimated following the Houseworth-Stahl interference escape model (also known as the gamma sprinkling or gamma escape model) [88] using the R package xoi v0.72 (Broman & Weber 2000). This model considers both Class I and Class II crossovers (where Class I crossovers are subject to interference, and Class II crossovers escape interference) and has been shown to be more robust than the gamma model, which considers only Class I crossovers [89]. For each focal ID, an estimate of ν and the fraction of crossovers escaping interference (*p*) was obtained by fitting the *fitStahl* function on crossover positions in all gametes from that individual. This resulted in a single estimate of ν and *p* for each focal ID. We used sex-specific cM positions rather than base pair (bp) positions, as interference acts on the length of the synaptonemal complex (SC) rather than sequence length. Recombination rates are highly correlated with SC length, and the relationship between SC length and base pair length varies between the sexes and along the genome in mammals [30,87].

### Estimation of heritabilities and genetic correlations

Additive genetic variances were estimated for crossover phenotype within each sex separately using a restricted maximum-likelihood “animal model” approach [88] in ASReml-R v4 [89]. The random effect structure included the additive genetic effect (fit with the inverse of the relationship matrix calculated from the pedigree) and the permanent environment effect (the identity of the focal ID) [90]. As crossover interference did not have repeated measures, the permanent environment was not fitted. The narrow sense heritability, h^2^, was estimated as the fraction of the total phenotypic variance V_P_ that was explained by additive genetic variance V_A_. The additive genetic correlation, r_A_, was estimated using bivariate animal models between all crossover phenotypes within each sex (cross-phenotype correlations), and for each crossover phenotype between the sexes (cross-sex correlations).

### Genome wide association studies (GWAS)

The association between 524,587 imputed SNPs and each crossover phenotype within each sex was tested using the mixed linear model leaving-one-chromosome-out (MLM LOCO) model implemented in GCTA version 1.93.2 beta Linux [91]. This option performs an association analysis fitting a model where the chromosome on which the candidate SNP is located is left out when calculating the genomic relationship matrix. The model was fit as follows:

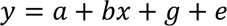

where *y* is the crossover phenotype, *a* is the mean term, *b* is the additive fixed effect of the SNP currently tested for association, *x* is the genotype dosage (0,1,2) and *g* is the accumulated effect of all markers except the markers on the chromosome where the SNP currently tested for association is located. The genome-wide significance level was determined using a Bonferroni correction, with the threshold of α = 0.05 set at P = 9.53×10^-8^. Candidate loci in significant regions were determined using biomaRt v2.60.0 in R v4.4.0 relative to Ensembl Release 112 [92] to extract Gene Ontology terms for loci within 500kb of the top associated SNPs. Gene names, GO names and GO descriptions were screened for the following text strings using the *grep* function: *meio*, *recombin*, *crossover*, *chromat*, *synapto*, *synapsis*, *gamet*, *double strand break*, *kinetoch*, *cohesin*, *histone*, *nucleosome*, and *spindle*. These flags were chosen to identify genes potentially associated with meiosis, synapsis, chromosome and chromatin structure, and gametogenesis. Selected candidates were screened for functions that matched the strings but not associated with meiosis (e.g. “synaptosome”). We then classified genes with GO terms matching the strings *meio*, *recombination*, *crossover*, *synaptonemal complex*, and *cohesin* as direct candidates, with the remainder classified as indirect candidates (Table S3).

## Acknowledgements

We thank Norsvin and Topigs Norsvin for providing access to the data for this study. Most analyses in this study have been executed on the Orion cluster at NMBU (https://orion.nmbu.no). We thank John McAuley, Roman Stetsenko, Bernard de Massy, Chloé Girard, Yukiko Imai and Sally Otto for discussions on the analysis and interpretation of the study. SEJ was supported by a Royal Society University Research Fellowship (UF150448 and URF/R/211008), with CB supported by Royal Society Enhanced Research Expenses (RF/ERE/231131 and RF/ERE/210071).

