## Supplementary material for "Individual variation in meiotic crossover positioning, rate and interference are associated with distinct genetic processes in domestic pigs"

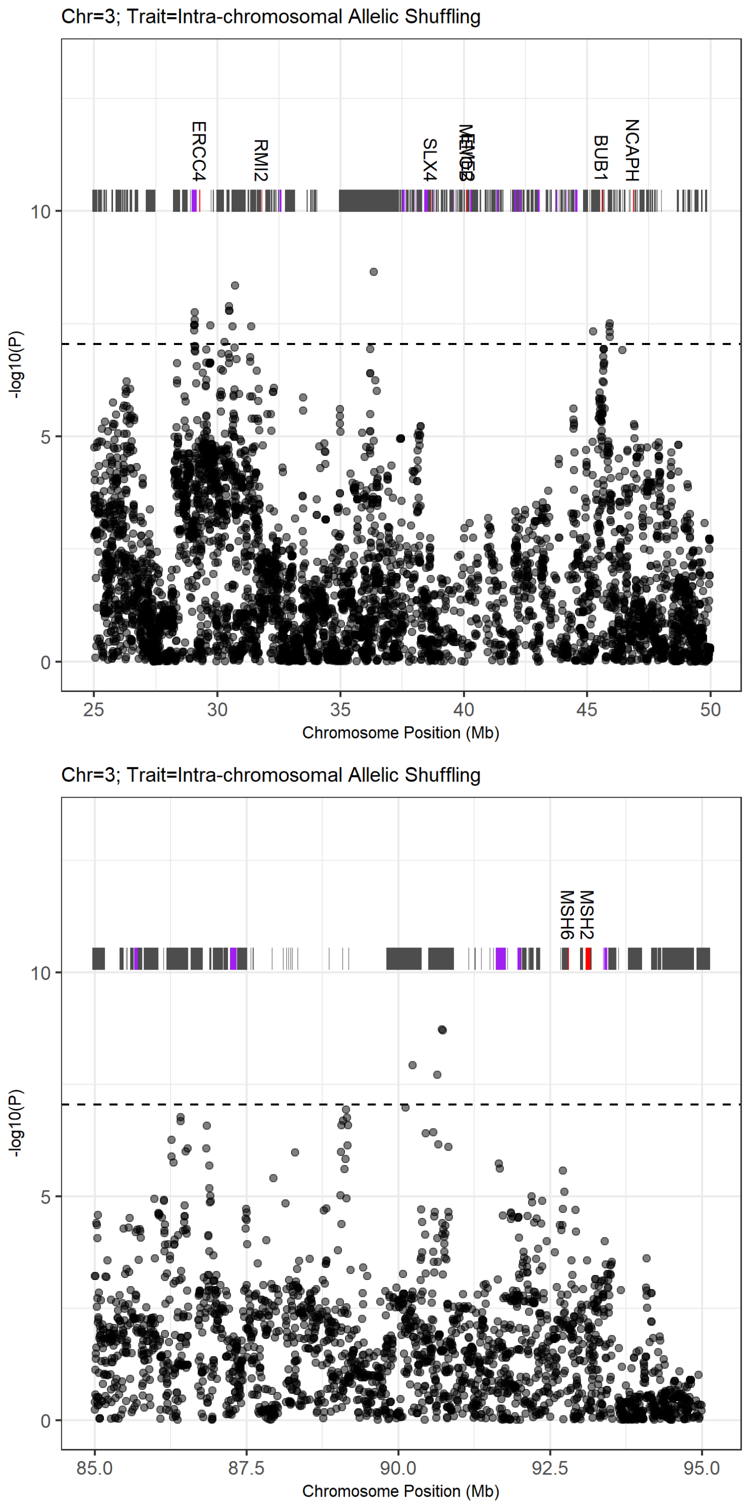


**Figure S1: Manhattan plots of genome regions significantly associated with crossover phenotypes**. Full details are provided in Table 1 of the main text. Plot titles provide information on the specific chromosome and the focal trait. The dashed line indicates the genome-wide significance threshold at $\alpha = 0.05$. Gene positions are indicated above the points and are coloured by their associations with meiotic processes. Dark grey = not associated, purple = indirectly associated, red = directly associated. Gene names are also shown for directly associated genes. All gene descriptions are provided in Table S3. Plots are continued on the following pages.

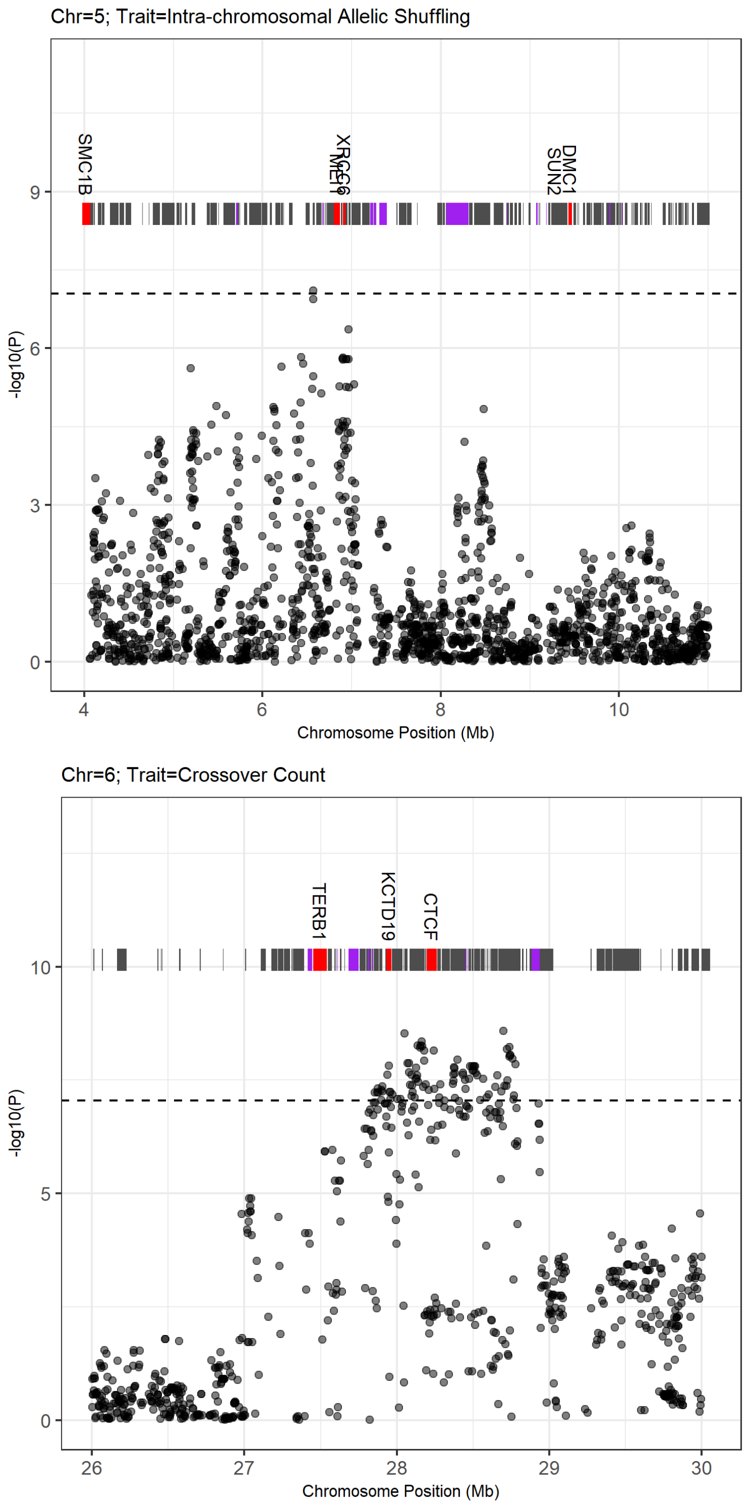


**Figure S1: Manhattan plots of genome regions significantly associated with crossover phenotypes.** ***(continued)***

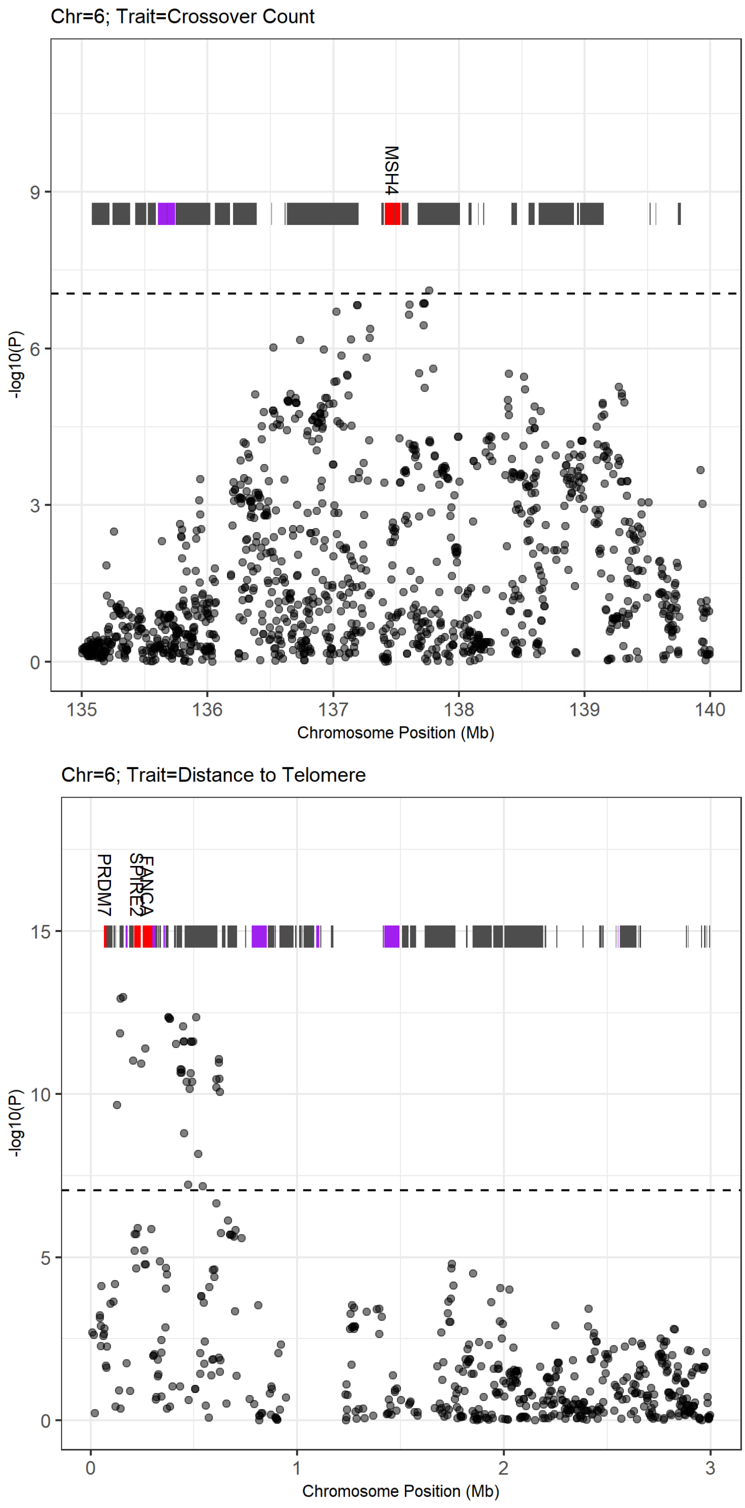


**Figure S1: Manhattan plots of genome regions significantly associated with crossover phenotypes. *(continued)***

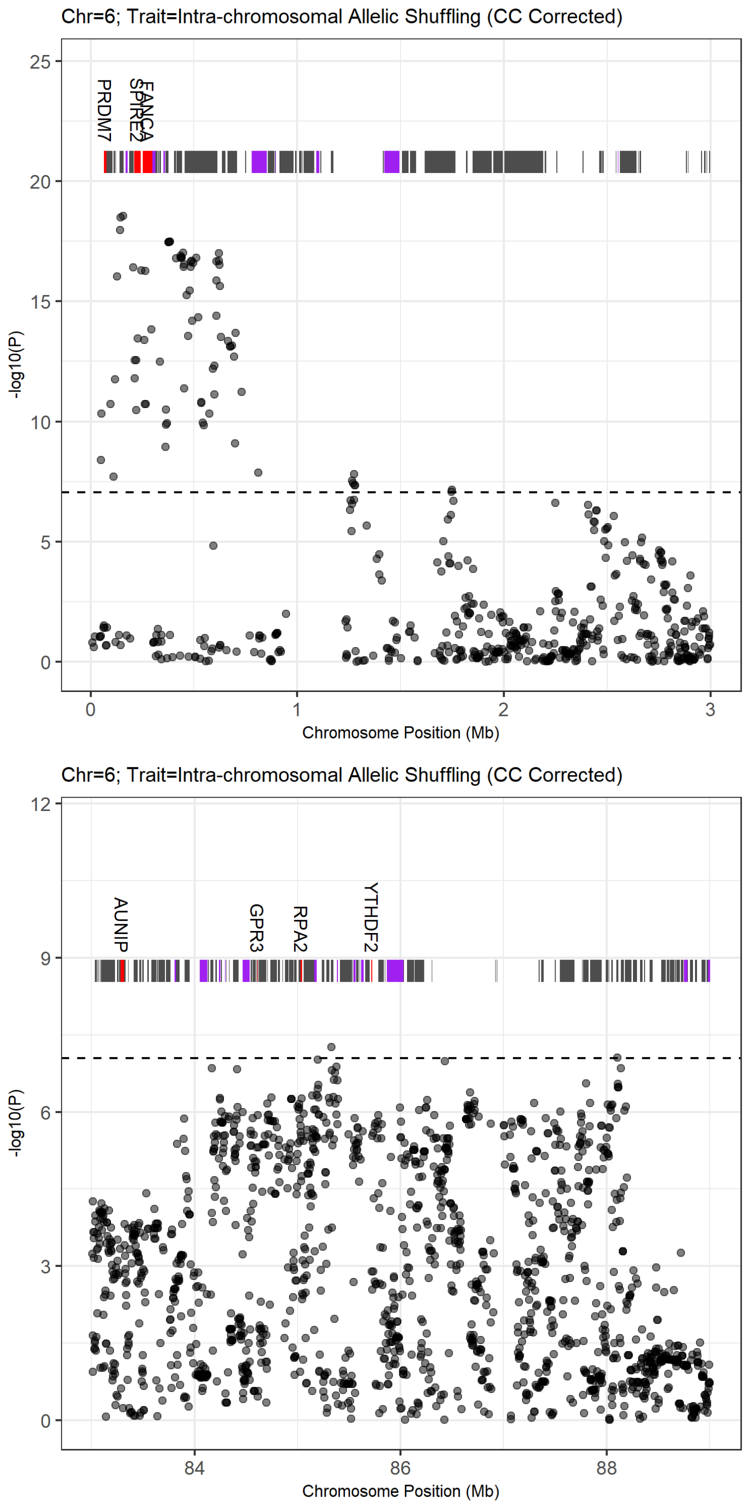


**Figure S1: Manhattan plots of genome regions significantly associated with crossover phenotypes.** ***(continued)***

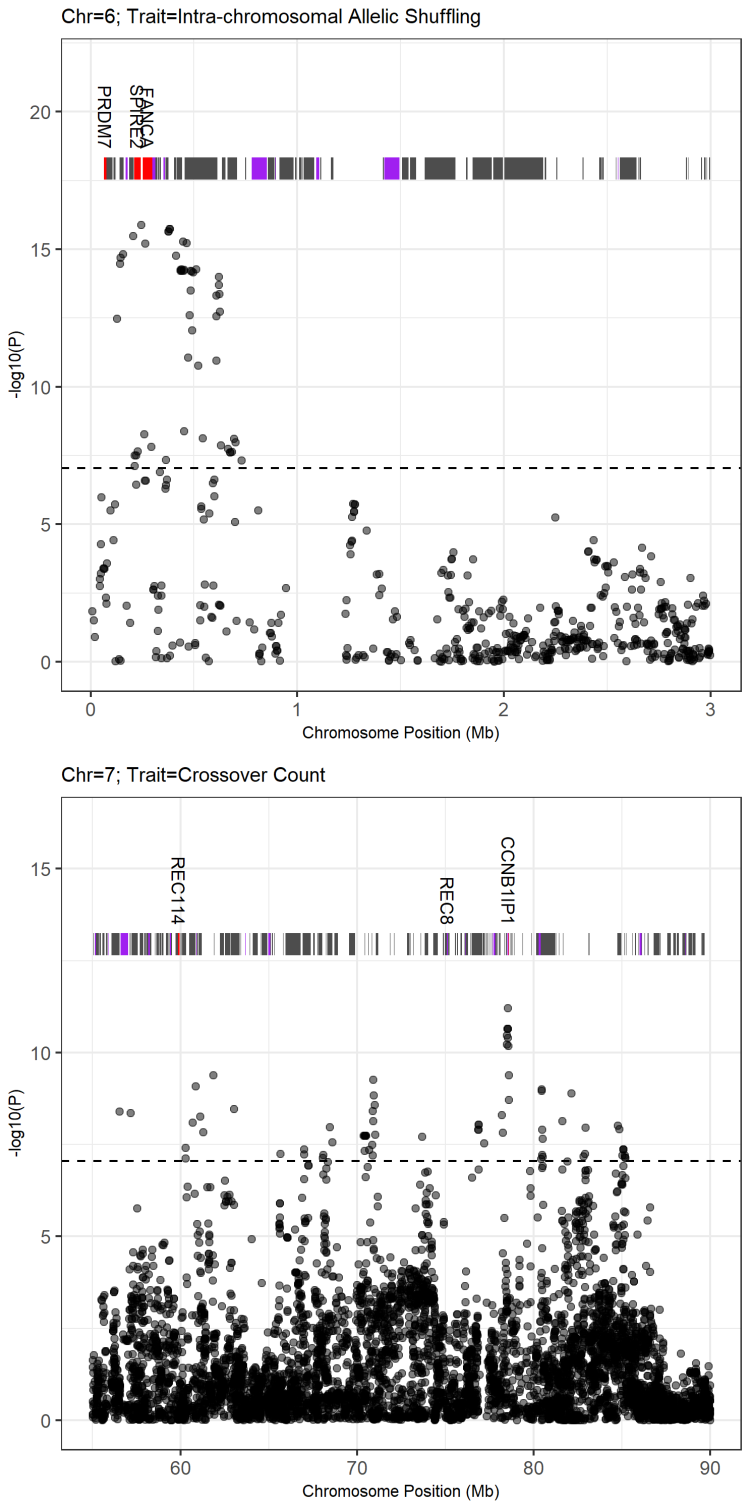


**Figure S1: Manhattan plots of genome regions significantly associated with crossover phenotypes. *(continued)***

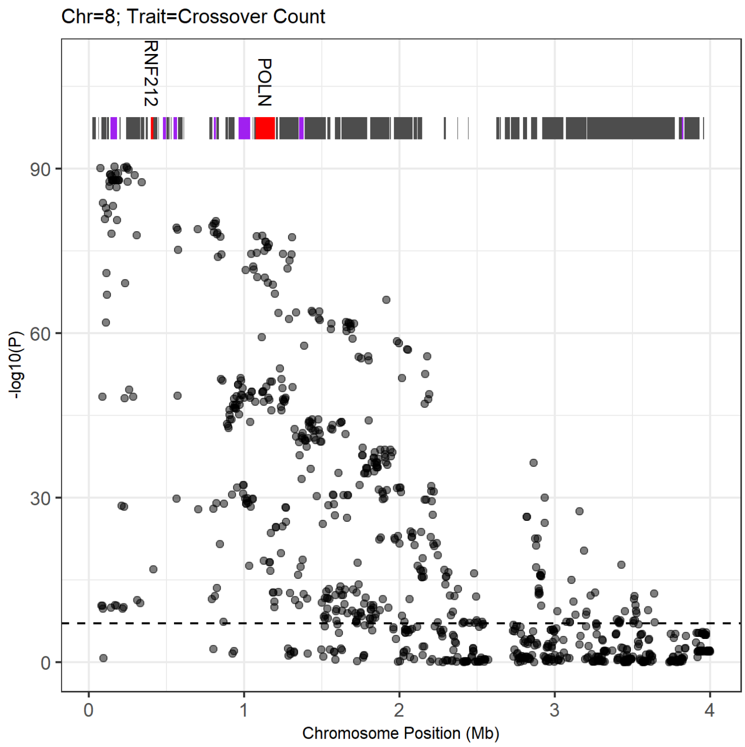


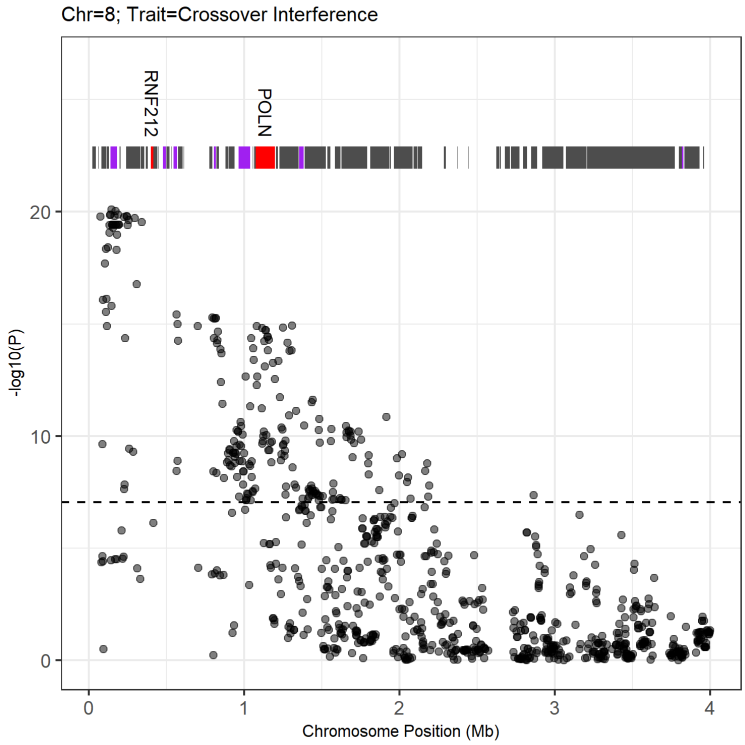


**Figure S1: Manhattan plots of genome regions significantly associated with crossover phenotypes. *(continued)***

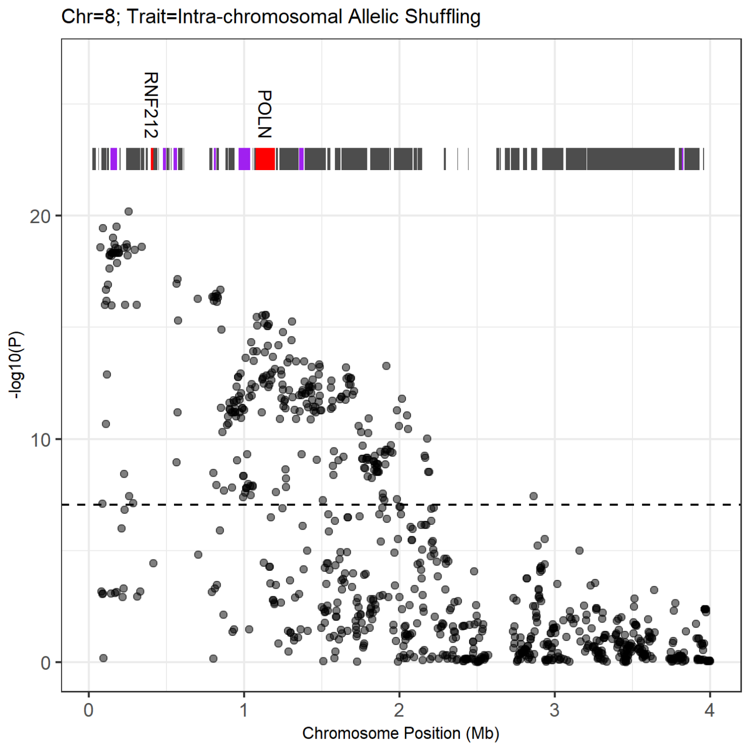


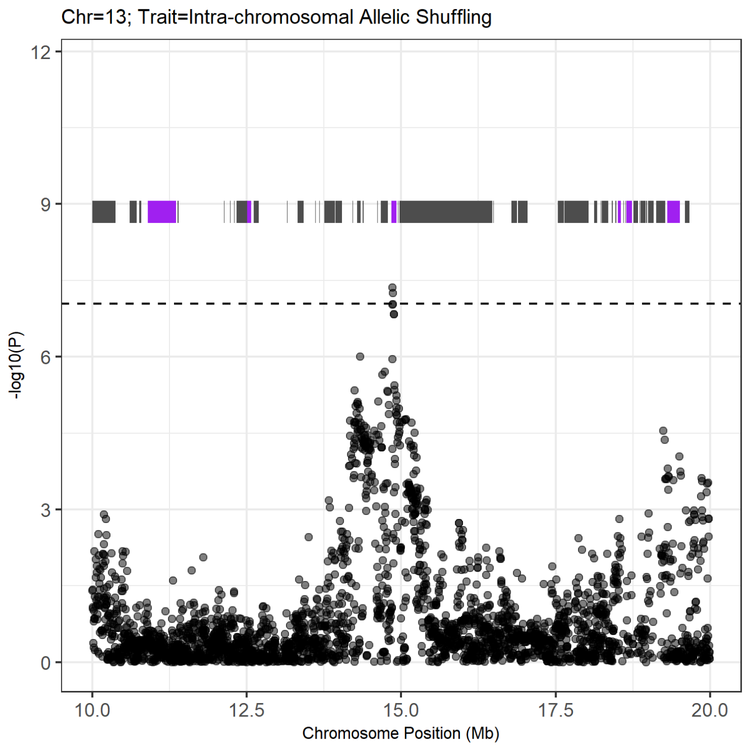


**Figure S1: Manhattan plots of genome regions significantly associated with crossover phenotypes. *(continued)***

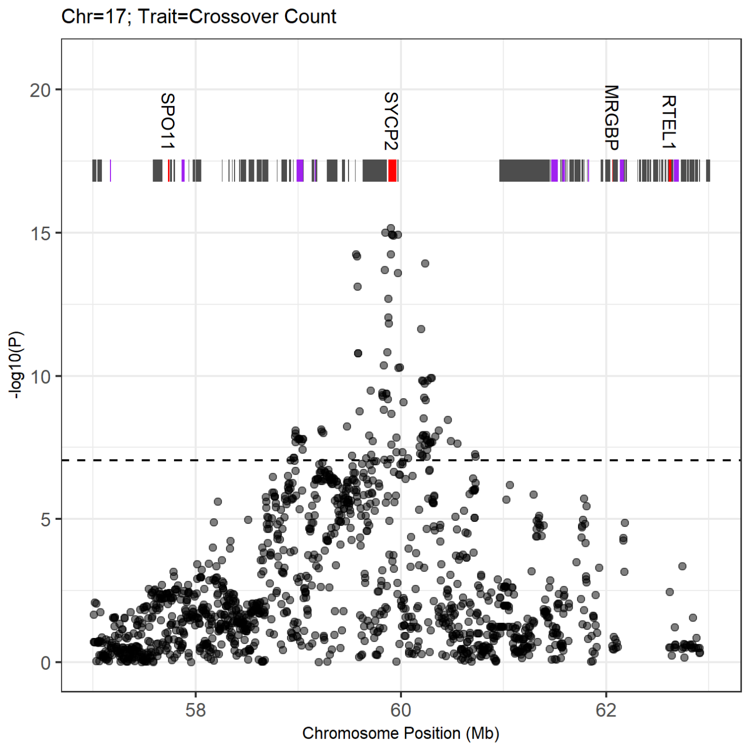


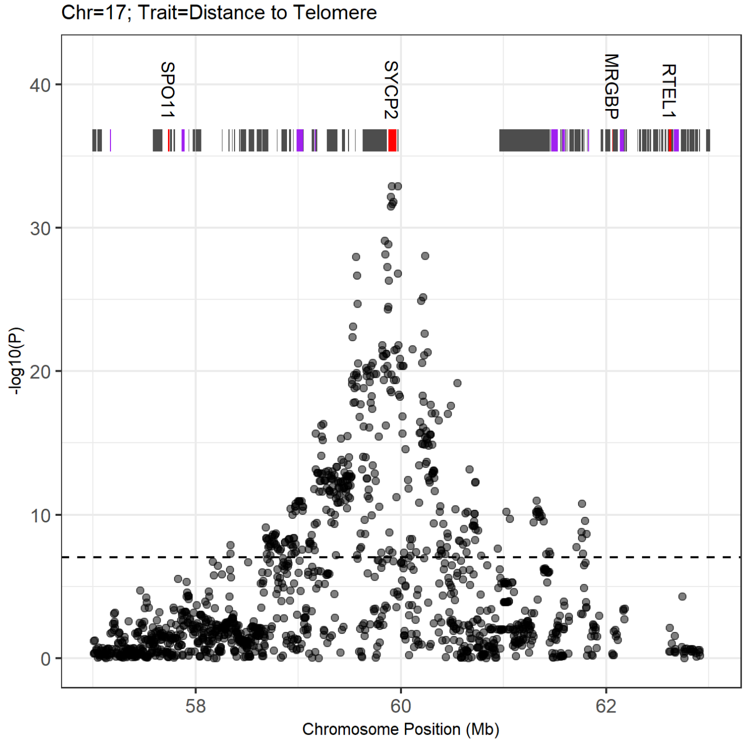


**Figure S1: Manhattan plots of genome regions significantly associated with crossover phenotypes. *(continued)***

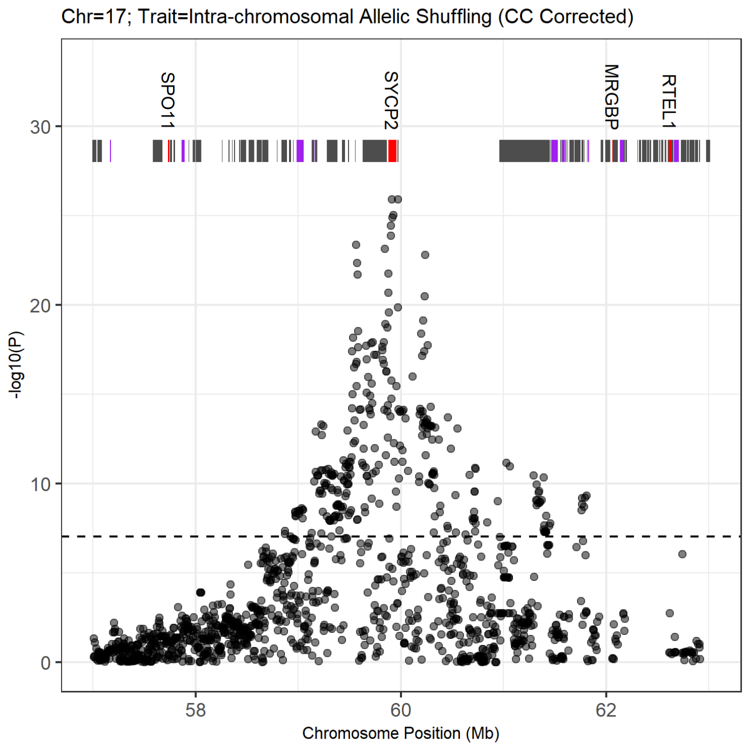


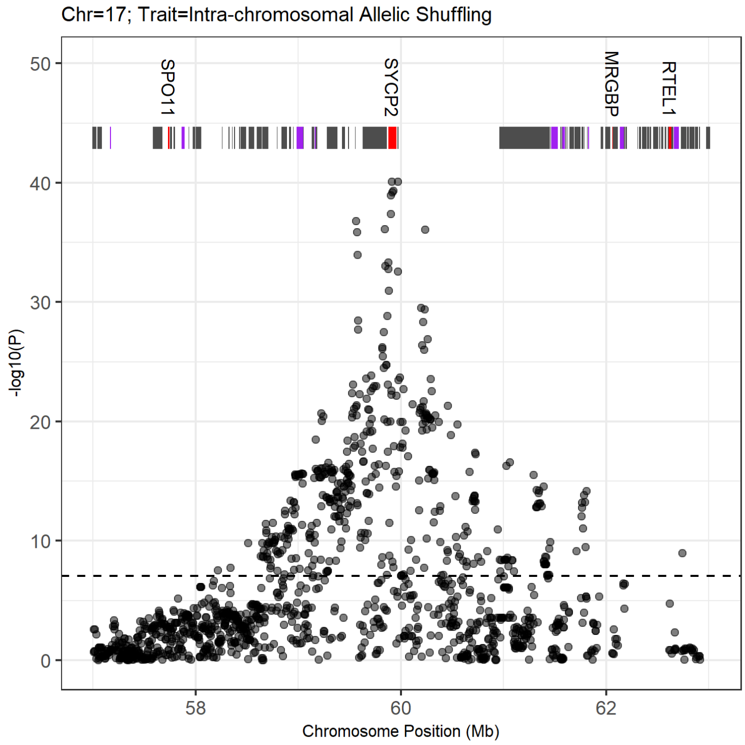


**Figure S1: Manhattan plots of genome regions significantly associated with crossover phenotypes.** ***(continued)***

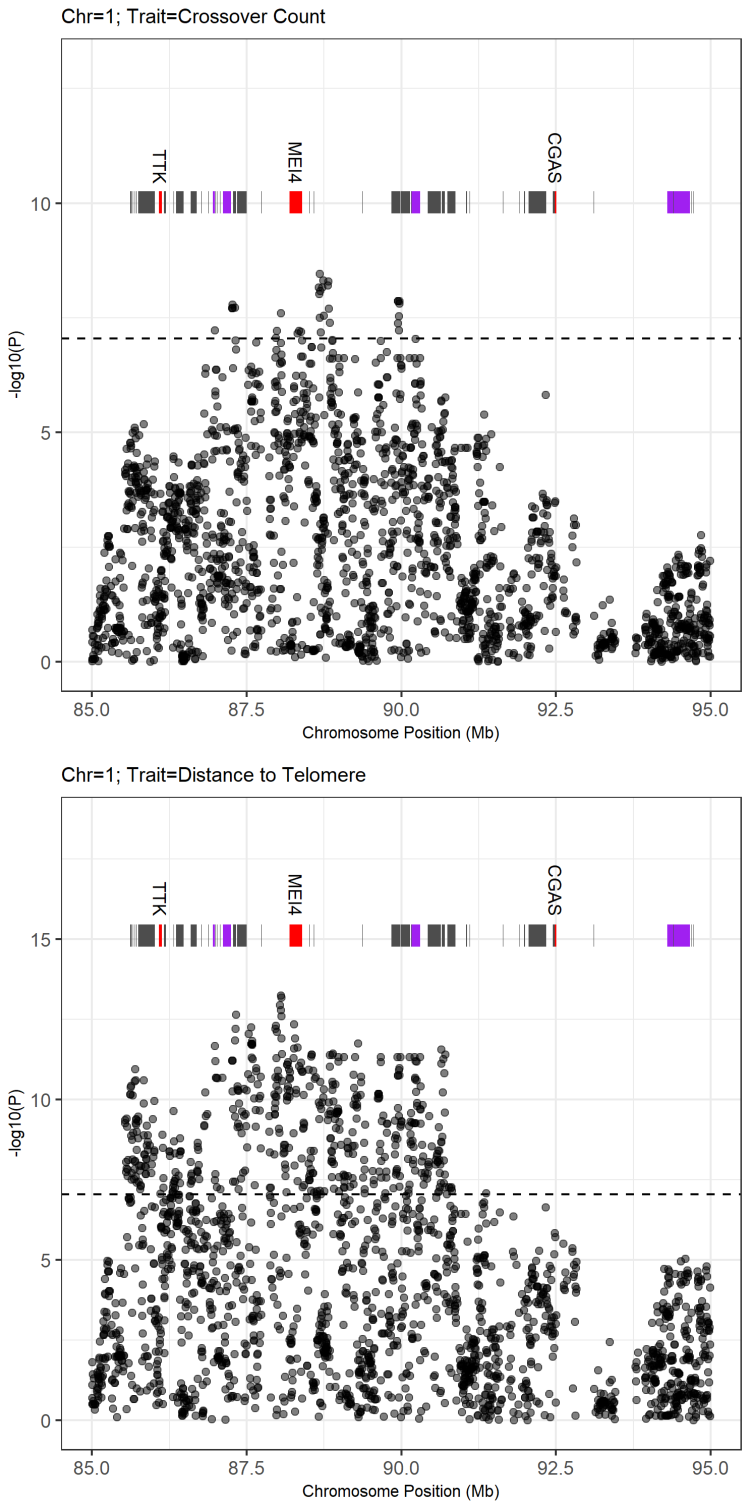


**Figure S1: Manhattan plots of genome regions significantly associated with crossover phenotypes.** ***(continued)***

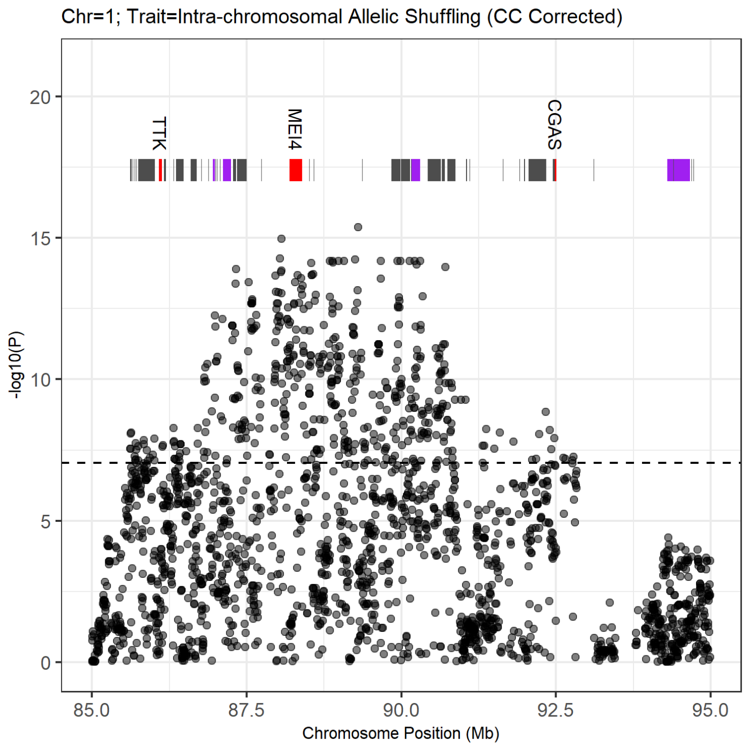


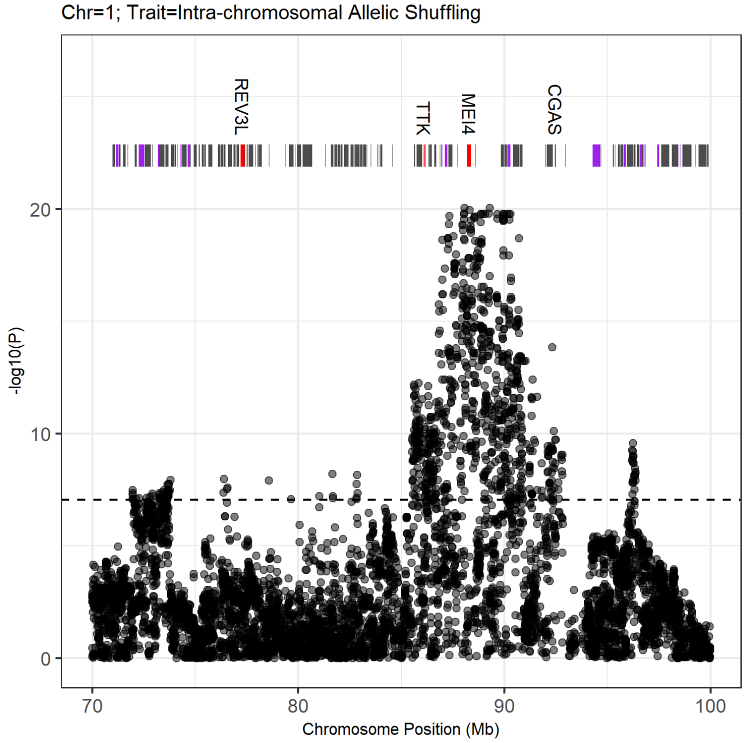


**Figure S1: Manhattan plots of genome regions significantly associated with crossover phenotypes.** ***(continued)***

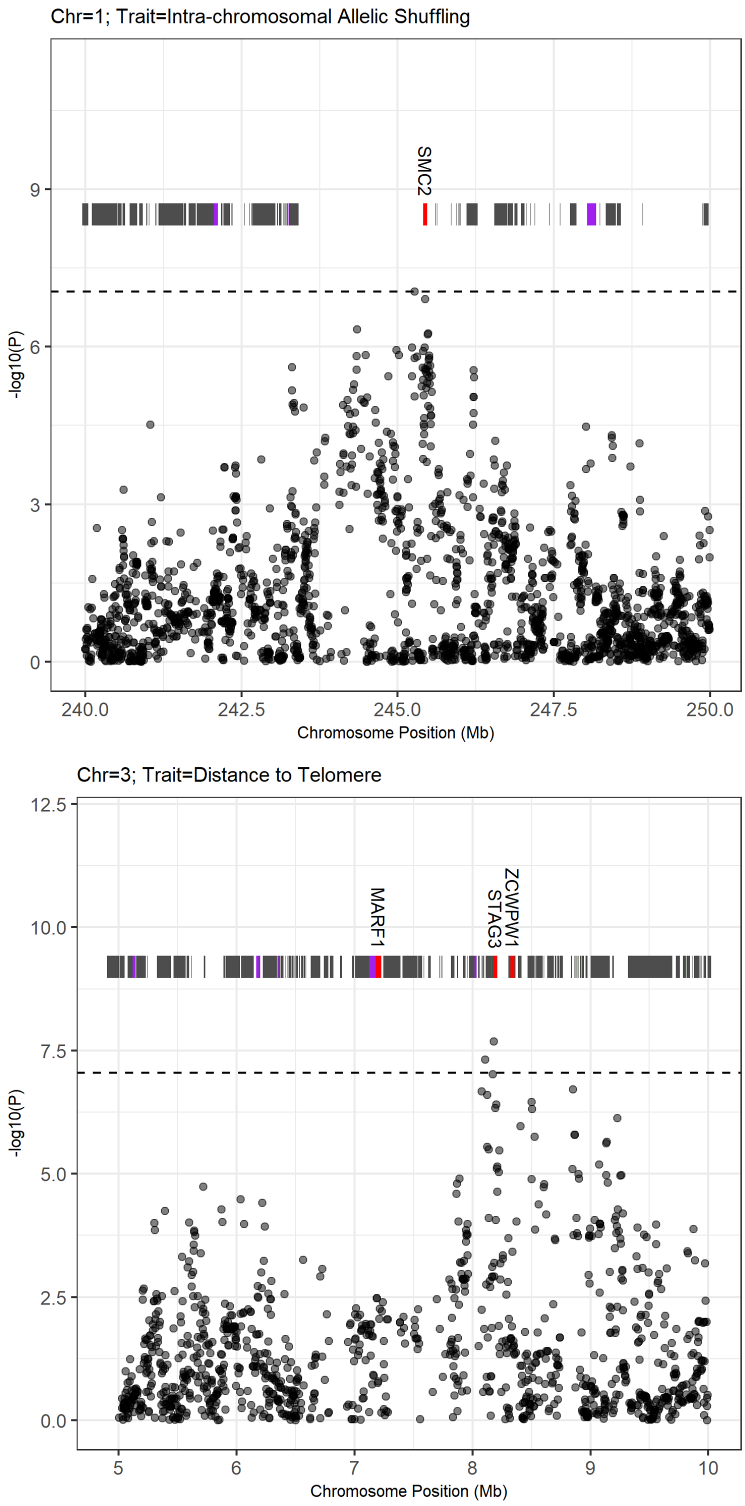


**Figure S1: Manhattan plots of genome regions significantly associated with crossover phenotypes.** ***(continued)***

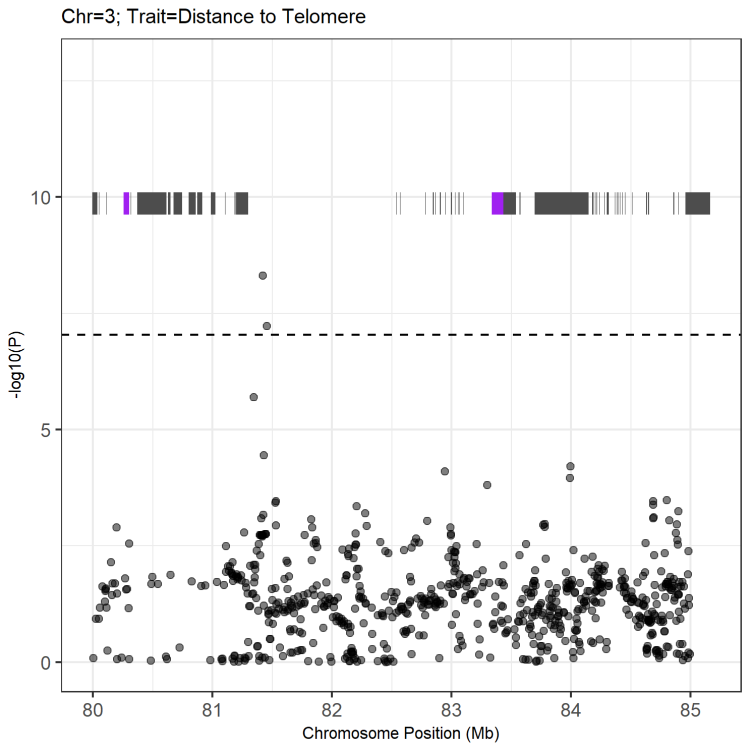


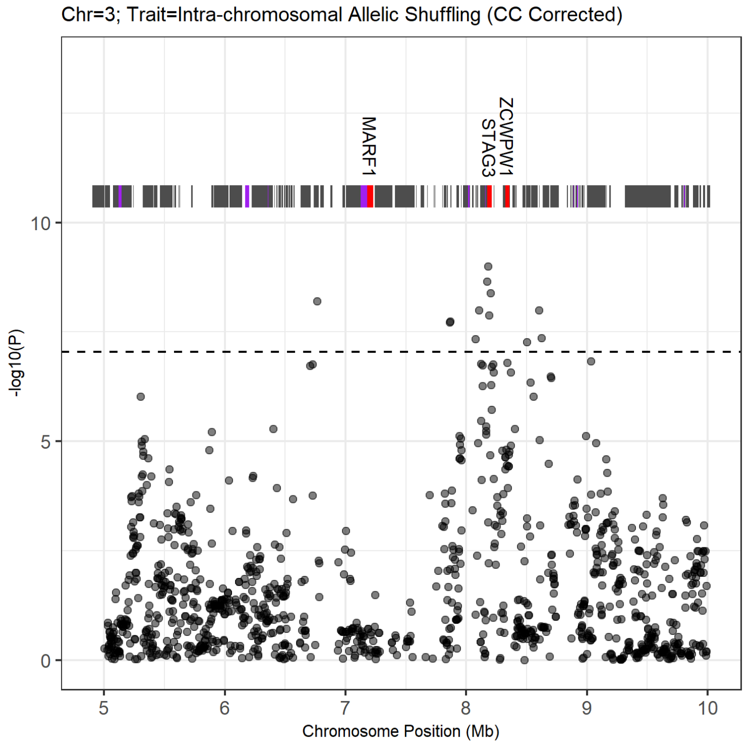


**Figure S1: Manhattan plots of genome regions significantly associated with crossover phenotypes. *(continued)***

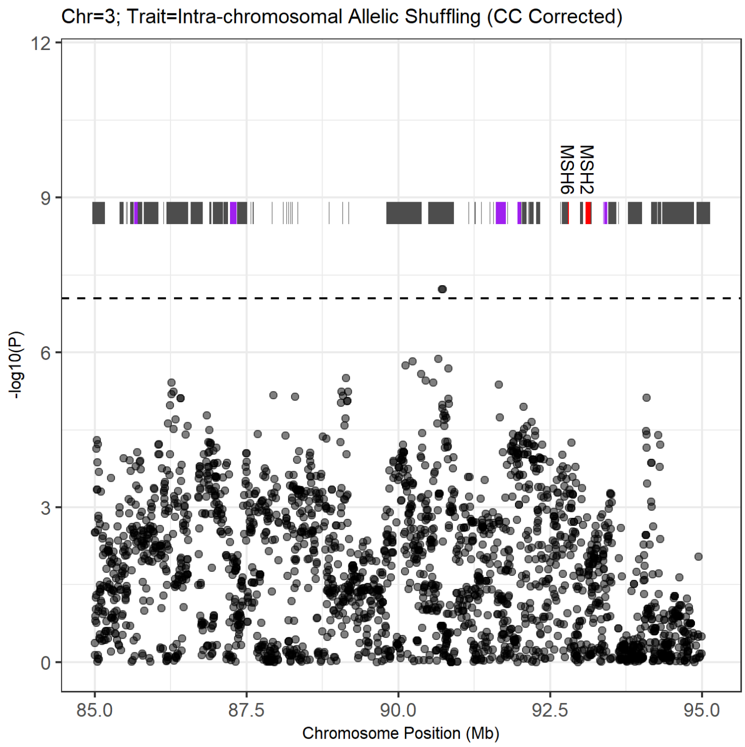


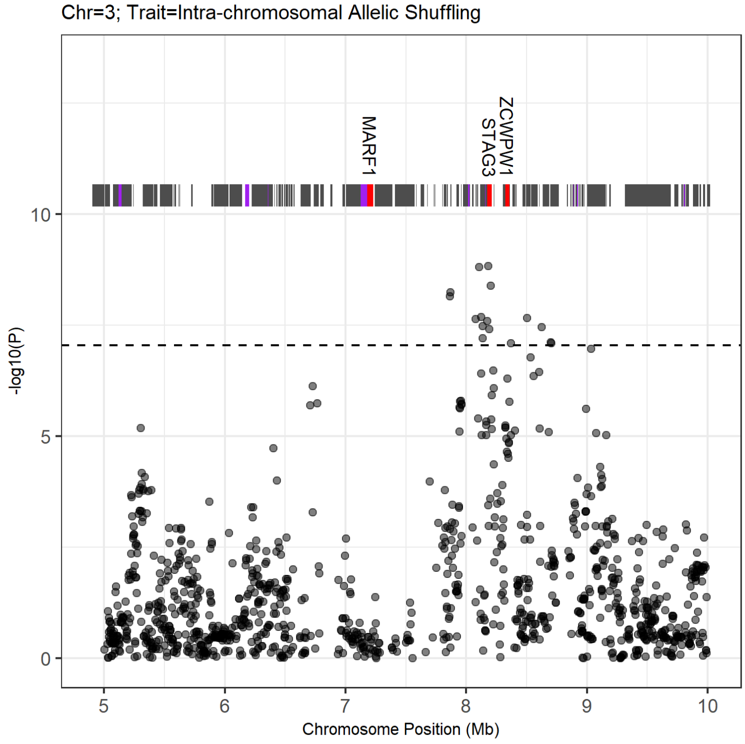


**Figure S1: Manhattan plots of genome regions significantly associated with crossover phenotypes.** ***(continued)***

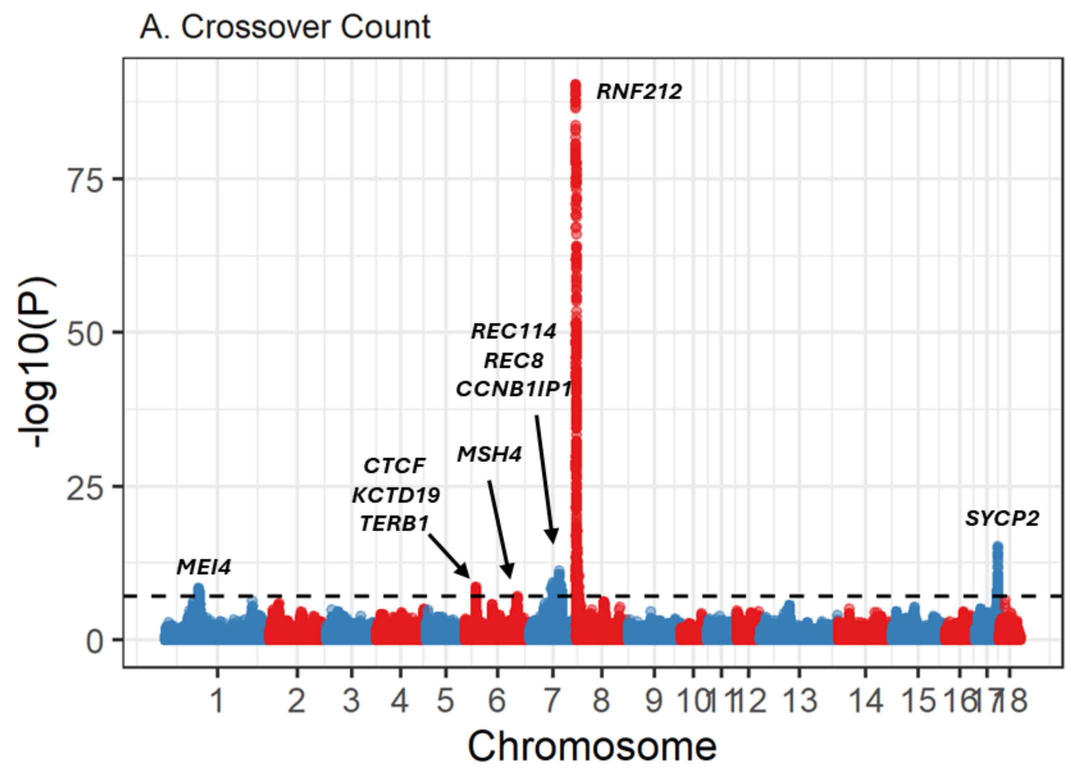


**Figure S2: Manhattan plot of genome regions significantly associated with female crossover count**. Associations are displayed at 554,768 SNPs. Sample sizes are provided in Table 1. Gene names above significant peaks indicate candidate loci. The dashed line indicates the genome-wide significance threshold at α= 0.05. Association statistics have been corrected with the genomic control parameter λ. Information on significant loci and candidate genes are provided in Table 2 and Tables S1-3.

**Table S1. Significant SNP loci for a GWAS on crossover phenotypes in female pigs.** All associations passed the Bonferroni significance threshold of α = 0.05. Columns are defined as follows. TRAIT = crossover phenotype CHR = chromosome, SNP = the SNP identifier, BP = SNP position in base pairs, A1 and A2 = alleles 1 and 2, AF1 = frequency of allele 1, B = effect size, SE = standard error of the effect size, P = significance value.

[Table_S1_All_Significant_SNPs.txt]

**Table S2: Information and positions of direct candidate genes occurring at significantly associated regions**. Full information on genes associated with which traits is provided in Table 1 of the main text.

| Gene Name | Chr | Gene Start | Gene Stop | Strand | Full Gene Name |
|---|---|---|---|---|---|
| *MEI4* | 1 | 88,189,903 | 88,399,137 | -1 | meiotic double-stranded break formation protein 4 |
| *SMC2* | 1 | 245,410,147 | 245,475,929 | 1 | structural maintenance of chromosomes 2 |
| *STAG3* | 3 | 8,175,915 | 8,211,889 | 1 | STAG3 cohesin complex component |
| *ZCWPW1* | 3 | 8,323,191 | 8,358,487 | -1 | zinc finger CW-type and PWWP domain containing 1 |
| *ERCC4* | 3 | 29,269,989 | 29,306,151 | -1 | ERCC excision repair 4, endonuclease catalytic subunit |
| *RMI2* | 3 | 31,808,015 | 31,814,843 | -1 | RecQ mediated genome instability 2 |
| *BUB1* | 3 | 45,585,226 | 45,624,543 | 1 | BUB1 mitotic checkpoint serine/threonine kinase |
| *NCAPH* | 3 | 46,852,392 | 46,893,013 | -1 | non-SMC condensin I complex subunit H |
| *MSH6* | 3 | 92,785,664 | 92,814,238 | -1 | mutS homolog 6 |
| *MSH2* | 3 | 93,081,219 | 93,163,585 | -1 | mutS homolog 2 |
| *MEI1* | 5 | 6,800,961 | 6,871,625 | -1 | meiotic double-stranded break formation protein 1 |
| *XRCC6* | 5 | 6,903,495 | 6,929,052 | -1 | X-ray repair cross complementing 6 |
| *PRDM7* (PRDM9)* | 6 | 64374 | 76394 | 1 | PR/SET domain 7. **NB.** This locus is likely to be *PRDM9* |
| *SPIRE2* | 6 | 211,487 | 243,561 | -1 | spire type actin nucleation factor 2 |
| *FANCA* | 6 | 253,207 | 300,152 | 1 | FA complementation group A |
| *TERB1* | 6 | 27,450,632 | 27,540,118 | -1 | telomere repeat binding bouquet formation protein 1 |
| *KCTD19* | 6 | 27,931,930 | 27,966,277 | -1 | potassium channel tetramerization domain containing 19 |
| *CTCF* | 6 | 28,195,647 | 28,261,009 | 1 | CCCTC-binding factor |
| *GPR3* | 6 | 84,610,107 | 84,613,202 | 1 | G protein-coupled receptor 3 |
| *RPA2* | 6 | 85,031,582 | 85,045,915 | -1 | replication protein A2 |
| *YTHDF2* | 6 | 85,716,983 | 85,726,473 | 1 | YTH N6-methyladenosine RNA binding protein F2 |
| *MSH4* | 6 | 137,410,691 | 137,534,233 | -1 | Rab geranylgeranyltransferase subunit beta |
| *REC114* | 7 | 59,838,480 | 59,941,418 | -1 | REC114 meiotic recombination protein |
| *REC8* | 7 | 75,118,744 | 75,125,537 | -1 | REC8 meiotic recombination protein |
| *CCNB1IP1* | 7 | 78,564,122 | 78,575,462 | 1 | cyclin B1 interacting protein 1 |
| *RNF212* | 8 | 399,715 | 413,896 | -1 | ring finger protein 212 |
| *ZCWPW2* | 13 | 14,841,319 | 14,926,248 | 1 | zinc finger CW-type and PWWP domain containing 2 |
| *SYCP2* | 17 | 59,875,993 | 59,955,209 | -1 | synaptonemal complex protein 2 |

**Table S3. Gene Ontology (GO) terms for genes occurring in regions significantly associated with crossover phenotypes that are associated with meiotic and related processes.** These are gene descriptions and/or GO terms that matched the strings "meio", "recombin", "crossover", "chromat", "synapto", "synapsis", "gamet", "double strand break", "kinetoch", "cohesin", "histone", "nucleosome", and "spindle". All terms that were associated with the strings “meio”, “recombination”, “crossover”, “synaptonemal complex”, and “cohesin” were flagged as direct candidates, with the remainder as indirect candidates. Columns are defined as follows: ensembl_gene_id = the Ensembl gene identifier, external_gene_name = gene name, chromosome_name = chromosome, start_position = gene start position, end_position = gene end position, strand = strand orientation, description = gene full name, go_id = GO term, Direct.Candidate.Flag = flagged as a direct candidate, name_1006 = GO term name, definition_1006 = GO term definition.

[Table_S3_Candidate_GO_Terms.txt]
